# Coral demographic performances in New Caledonia, a video transect approach to operationalize imagery-based investigation of population and community dynamics

**DOI:** 10.1101/2023.05.12.540552

**Authors:** Mohsen Kayal, Eva Mevrel, Jane Ballard

## Abstract

Demographic studies that quantify species performances for survival, growth, and reproduction are powerful means to understand and predict how species and communities respond to environmental change through the characterization of population dynamics and sources of demographic bottlenecks. However, demographic studies require fine-scale surveys of populations in the field, and are often too effort-intensive to be replicable at large scale and in the long-term. To surpass this obstacle, we developed a digital approach for extracting demographic data on species abundances, sizes, and positions within video-transects, facilitating back-from-the-field data acquisitions on population and community dynamics from video surveys. The approach is based on manual coral identification, size-measurements, and mapping along video-transects, mimicking what is traditionally performed in the field, thought it can be automated in the future with the deployment of artificial intelligence. We illustrate our approach with the characterizations of species demographic performances using surveys of a reef-building coral community in New Caledonia recorded with underwater cameras, therefore optimizing time spent in the field. The results provide quantitative measures of coral community composition and demographic performances as key ecological indicators of coral reef health, shed light on species life strategies and constraints to their demographics, and open paths for further quantitative investigations. Key findings include the diversity of species life strategies in terms of relative investment in survival, growth, and reproduction found among taxa dominating the coral community, indicating the diversity of demographic paths to ecological success and that several species have adapted mechanisms to prevail under limiting hydrodynamic environments. Our approach facilitates image-based extractions of demographic data, helping to accelerate empirical endeavors in ecology and ecosystem management.

**Author summary:** Sustainable ecosystem management requires comprehension of key ecological processes that affect species resilience. Accurate and reoccurring measurements of species helps us understand how they are responding to various environments and predict what might happen in the future. We developed a digital approach that mimics measurements traditionally performed in the field to measure species abundance, size, and distributions using video records of the ecosystems. This transition to imagery-based surveys helps researchers and managers acquire fine-scale ecological data while optimizing time spent in the field, particularly for studying remote and extreme environments where access is limited. We illustrate the application of our approach by characterizing the dynamics of a coral community in the vast tropical reef system of New Caledonia, where such evaluations of demographic processes controlling coral resilience are inexistent but necessary.

## Introduction

As ecosystems weather growing environmental changes and human impacts, performing ecological diagnostics is increasingly crucial to anticipate declines and identify solutions supporting ecosystem resilience (Halpern et al. 2019; Weiskopf et al. 2020; Condie et al. 2021). Demographic studies that quantify species dynamics and performances in key life cycle processes, such as survival, growth, and reproduction, are powerful tools for characterizing life-strategies, influential regulatory mechanisms, and community trajectory predictions (Ellner et al. 2016). However, the high level of effort necessary to perform demographic surveys that track individual organisms over significant timeframes (e.g., multi-annual surveys) prevents a wider use of demographic approaches to inform management. We developed a digital approach for mapping and measuring individual organisms in fixed transects captured by video, replacing data collection tasks previously performed for long hours in the field, to characterize community structure at each observation (species composition, abundance, and size-distribution), and demographic performance between consecutive surveys (survival, growth, reproduction, and migration rates; Kayal et al. 2015). We describe the application of our approach on a reef building coral community in New Caledonia.

Coral reefs are central to marine biodiversity and human well-being but declining due to increasingly altered coastal environments associated with coastal development, pollution, fishing, and climate change (Darling et al. 2019; Hughes et al. 2019; Halpern et al. 2021). The declines in coral community abundance, composition, and size are major threats for tropical marine biodiversity and the societies that depend on these ecosystems for services such as food, shelter from storm waves, and economic and social fulfillment (Hoegh-Guldberg et al. 2019; Eddy et al. 2021; Carlot et al. 2023). Researchers and managers currently use fine-scale surveys of coral populations to investigate the demographic mechanisms controlling coral reef resilience and predict future trajectories (Kayal et al. 2018; Madin et al. 2014; Riegl et al. 2018; Carlot et al. 2021). However, these efforts lag far behind the challenge of coral reef conservation in the twenty-first century (Halpern et al. 2019), and are often restricted to only a few eminent sites that benefit from high concentrations of scientific focus, leaving out most of the world’s coral reefs. Because species mapping and size measurements are traditionally performed by hand in the field, with only field notes being recorded, the shift to image-based data extractions and archiving significantly improves the efficiency of data collection and archival. This need is particularly acute for understanding remote and extreme environments, such as underwater. While the development and accessibility of high-definition cameras have opened paths for increased imagery based approaches to ecosystem surveys, analytical tools for characterizing population and community dynamics remain restricted. We illustrate the application of our digital approach by characterizing the abundance, composition, size-distribution, and demographic performances of a coral community on the outer-reef of New Caledonia over one year, and describe benefits for ecological investigations into the future.

## Materials and methods

### Field survey and data extractions

We recorded six contiguous 5 m × 0.8 m (4 m^2^) video transects along a randomly selected representative 30 m stretch of the reef substrate at a permanent study site situated at a mid-range, 10 m water depth on the outer-reef slope of New Caledonia’s south-western barrier reef in March of 2021 and 2022 (Supplementary material S1 Figure). In part due to the relatively small and concentrated human population compared to the size of the coral system, New Caledonia reefs are among the most diverse and healthy in the world, and the mid-range depth on the outer-reef is where the highest taxonomic diversity is typically found (Fenner & Muir 2008; Andréfouët et al. 2009; Adjeroud et al. 2019; Kayal & Adjeroud 2022). Video transects were recorded on SCUBA (self-contained underwater breathing apparatus) using GoPro cameras facing downward (90°) towards the substrate at a distance of 50 cm from the transect tape. All individual reef-building corals (scleractinians and the calcifying hydrozoan *Millepora*) in the video transects were identified by genus and morphotype, measured in two dimensions (length, width), and the position of their centroid was mapped using X-Y coordinates (Fig. 1), an image-based adaptation of what is traditionally achieved in situ during long dives (Kayal et al. 2015, 2018). No major disturbance to the reef was recorded over the one-year span of the survey.

**Fig. 1.**
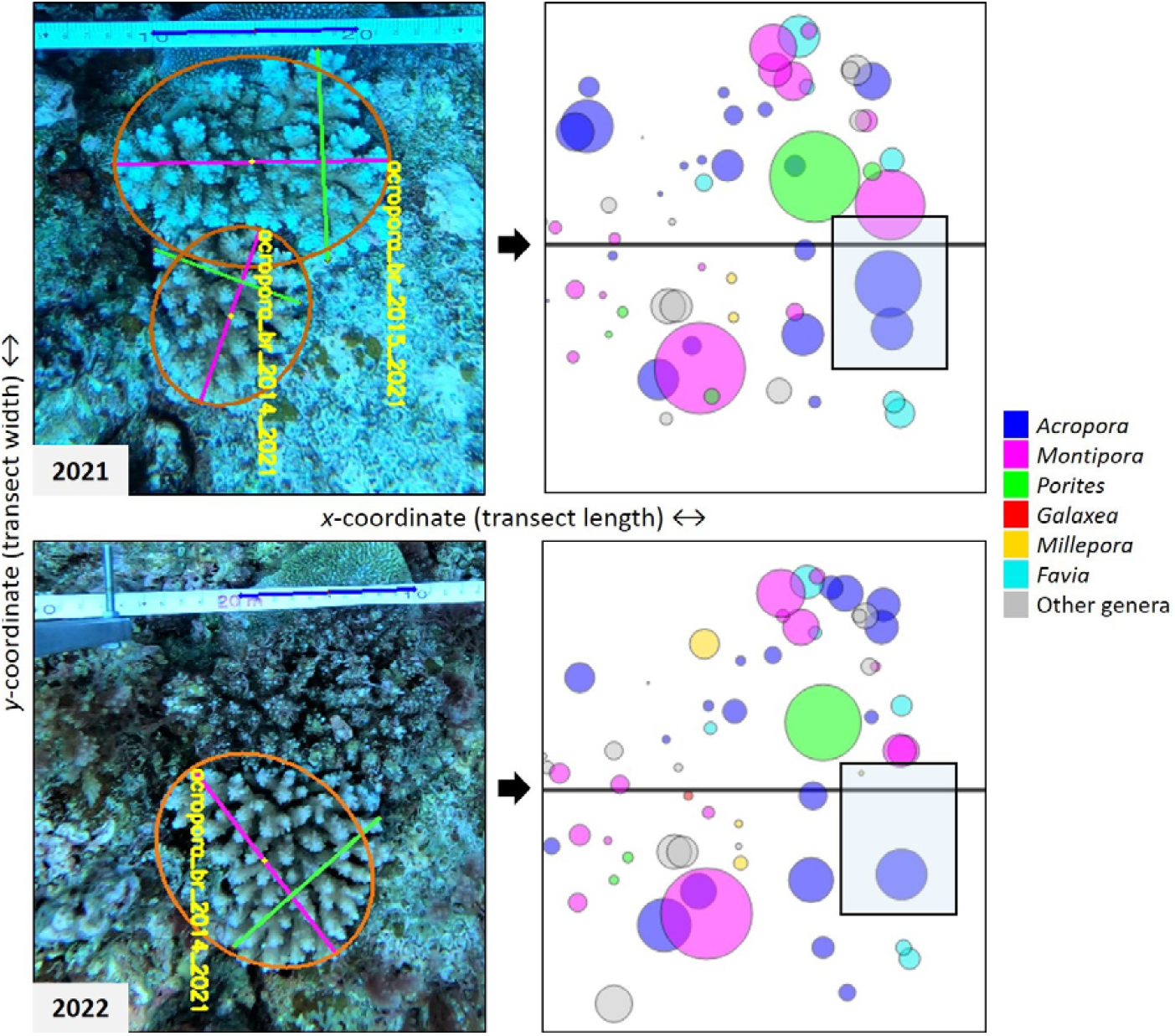
A portion of the video transects recorded in 2021 and 2022 illustrating our coral mapping and size measurement approach (pictures) and the resulting visualization (plots). Each coral is identified by genus and morphotype and given a unique identification code (yellow text) made of genus, morphotype, *x*-coordinate, and year of first observation. Colony size is estimated by measuring length (pink line) and width (green line) in pixels and converting to centimeters using a 10-cm reference distance along the transect tape (blue line). The mean coral diameter = (length+width)/2 is then referred to as coral size. The position of the centroid of each colony (yellow dot) is mapped along the length (*x*-coordinate) and width (*y*-coordinate) of the transect. Plots are visualizations of coral distribution (circle location), size (circle size), and composition (color code) along the transect. This example shows the dynamics of two branching *Acropora* colonies between 2021 and 2022, one growing and the other one dying.

Coral mapping and measurements are performed by extracting individual video frames, and measuring coral size and position relative to the graduated transect tape (Fig. 1). Only coral colonies entirely visible in the videos were considered, excluding those only partially visible. Information on coral taxonomic identity, morphotype, size, and position within transects are recorded in a data spreadsheet, enabling the characterization of coral community structure for each survey as well as population dynamics between surveys. Individual coral performances in terms of survival, growth, and recruitment are characterized as described in Kayal et al. (2015). New recruits were defined as small corals absent from the 2021 video transects and visible in 2022. Other coral dynamics (fragmentation, fission, fusion, and migration) were also quantified, but not considered in this study. Data extractions from the video transects were coded in Python complemented with OpenCv (https://opencv.org) and Tkinter (https://pythonbasics.org/tkinter/) libraries (S2 Appendix). Coral identification, mapping, and size-measurements were performed by the same observer.

### Data analysis

For each year (2021 and 2022), we characterized coral community abundance, composition, and size-distribution of the six dominant coral genera, for which we used individual coral dynamics between the two surveys to estimate annual demographic performances in survival, growth, and recruitment. Note that for coral survival and growth estimates, the sampling unit is the coral colony, not the transect. We accounted for size-dependent variation in our estimations of coral survival by relating survival probability to year *y*+1 to initial size at year *y* using generalized linear mixed-effect models accounting for random effects of longitudinal observations on individual colonies (Kayal et al. 2015). Similarly, size-dependent growth was characterized by relating final size at year *y*+1 to initial size at year *y*. Coral sizes were log-transformed to reduce data dispersion, and the models were checked for assumptions of residual homogeneity. For coral abundance and recruitment estimates, preliminary analysis showed similar results when considering the 30 m stretch of the reef recorded in video as six contiguous 5 m-long transects, versus dividing it into five 5 m-long transects spaced by 1 m gaps, and the former was retained. All colonies were used to estimate the size-distribution of coral populations. Modeling and graphing were performed in R (www.R-project.org) complemented with the NLME package (Pinheiro et al. 2023).

## Results

### Coral community structure

We identified, mapped, and measured 1,104 corals from 26 different genera over the two years of our survey (2021-2022); 894 corals were recorded in 2021 and 838 in 2022, with 628 colonies observed in both years (Fig. 2). The corals that were not observed both years consisted of new recruits in 2022 (84), corals that had died between the two years (213), and those situated in the shadows or on the edges of the video-transects and only visible in one of the survey years (157).

**Fig. 2.**
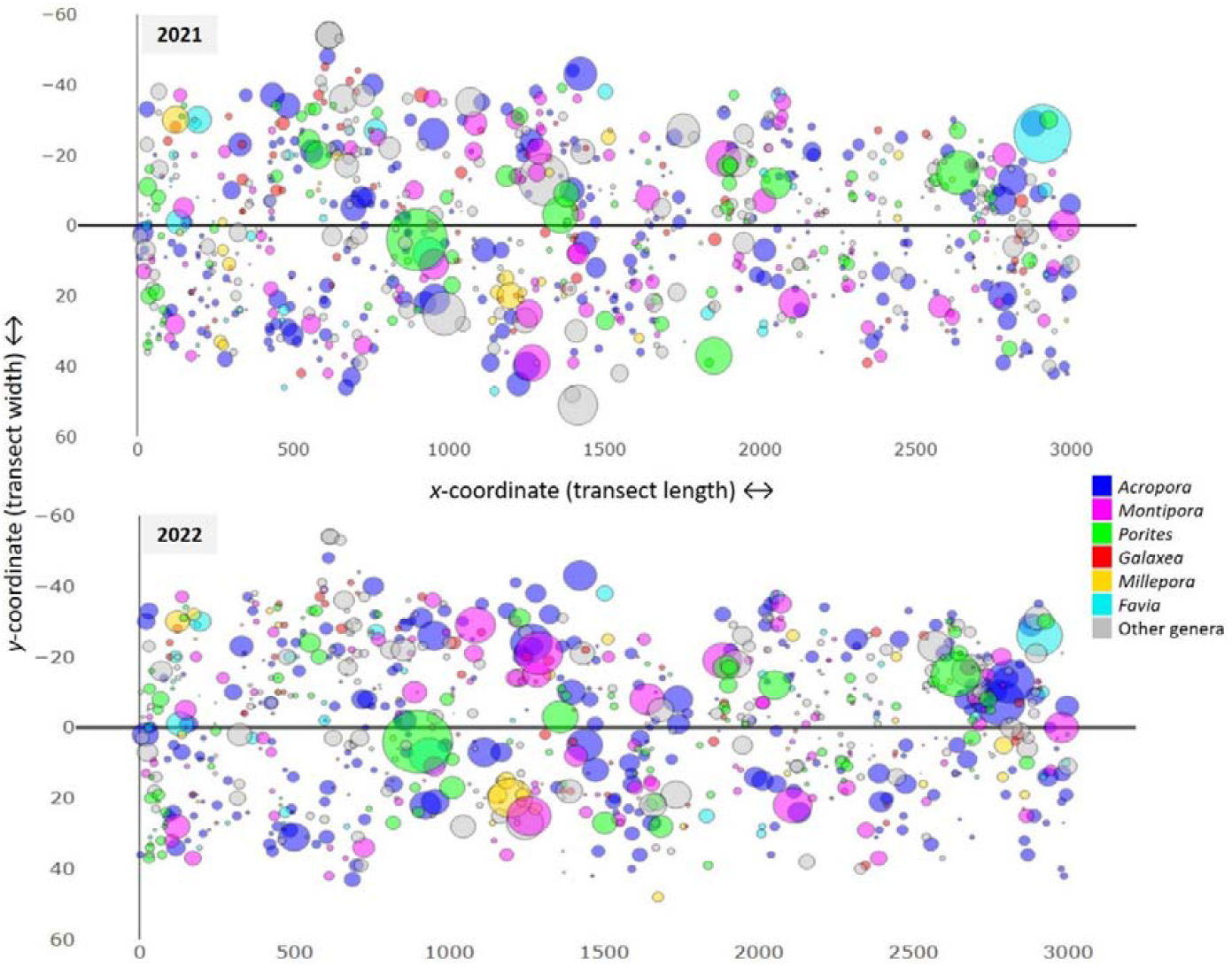
Coral spatial distribution, size, and taxonomic composition along the studied 30 m stretch of the reef substrate in 2021 and 2022. Each circle represents a coral with dimensions that are proportional to mean colony size. Color code distinguishes the six dominant genera.

Coral community abundance averaged 149 colonies ± 24 SD (standard deviation) per 4 m^2^ transect (equivalent to 37 corals per m^2^) in 2021, and 140 corals ± 26 SD per 4 m^2^ (35 corals per m^2^) in 2022. Six genera dominated the coral community with, over the two surveys, 30% of colonies belonging to *Acropora*, 14% *Montipora*, 12% *Porites*, 9% *Galaxea*, 6% *Millepora*, and 6% *Favia* (Fig. 3, S3 Table).

**Fig. 3.**
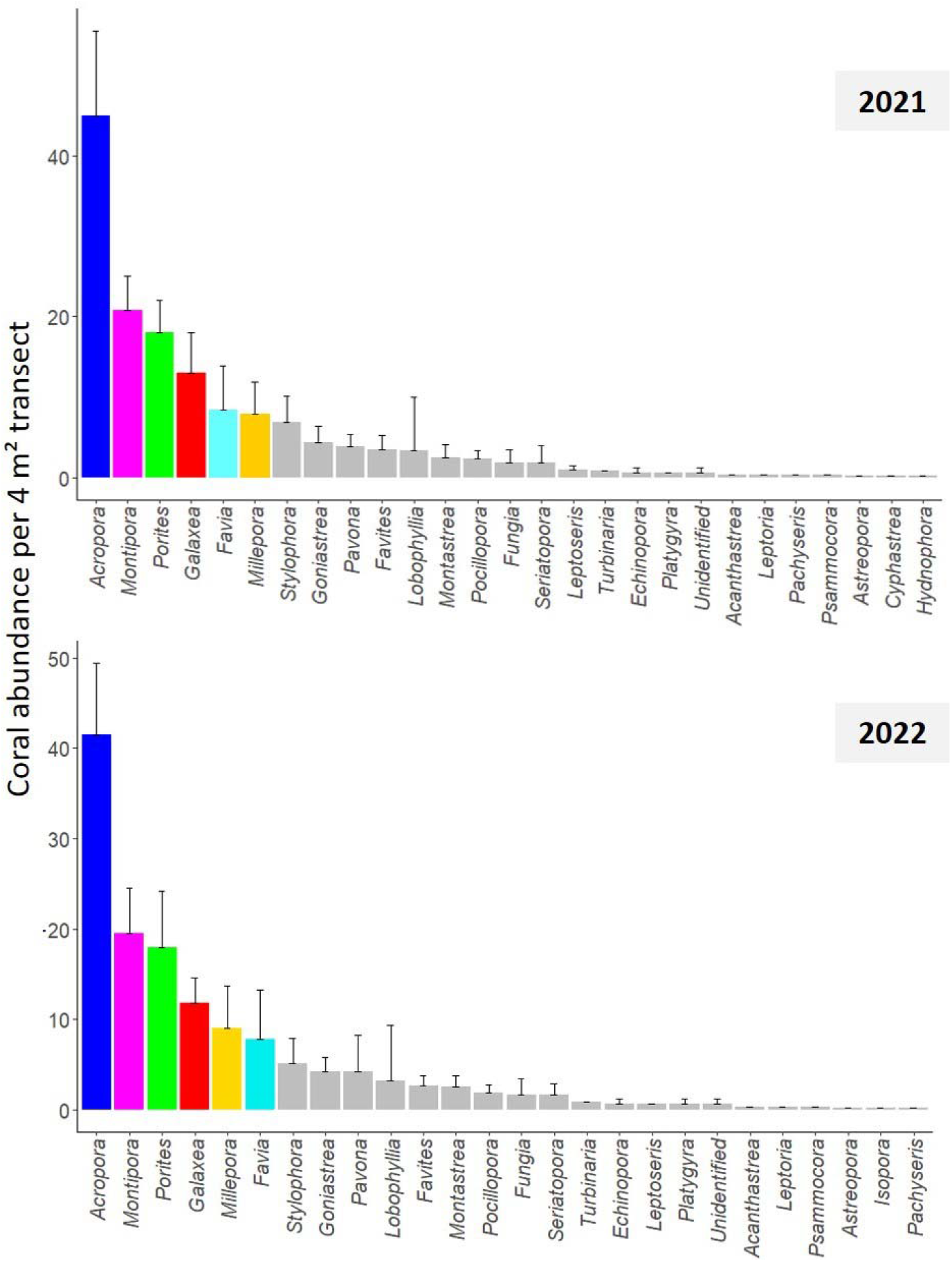
Coral population abundances (mean ± standard deviation) per 4 m^2^ transect by genus in 2021 and 2022. Color code distinguishes the six dominant genera. See S3 Table for details. We compared the size distributions of populations among the six dominant coral genera in 2021 and 2022 using Kolmogorov-Smirnov tests (S4 Table). Overall, *Acropora, Montipora*, and *Porites* populations were characterized by larger coral sizes, with mean colony diameters >5.6 cm (Fig. 4, S5 Figure). In contrast, *Galaxea, Favia*, and *Millepora* populations were characterized by higher proportions of small colonies, with mean diameters <5.3 cm.

**Fig. 4.**
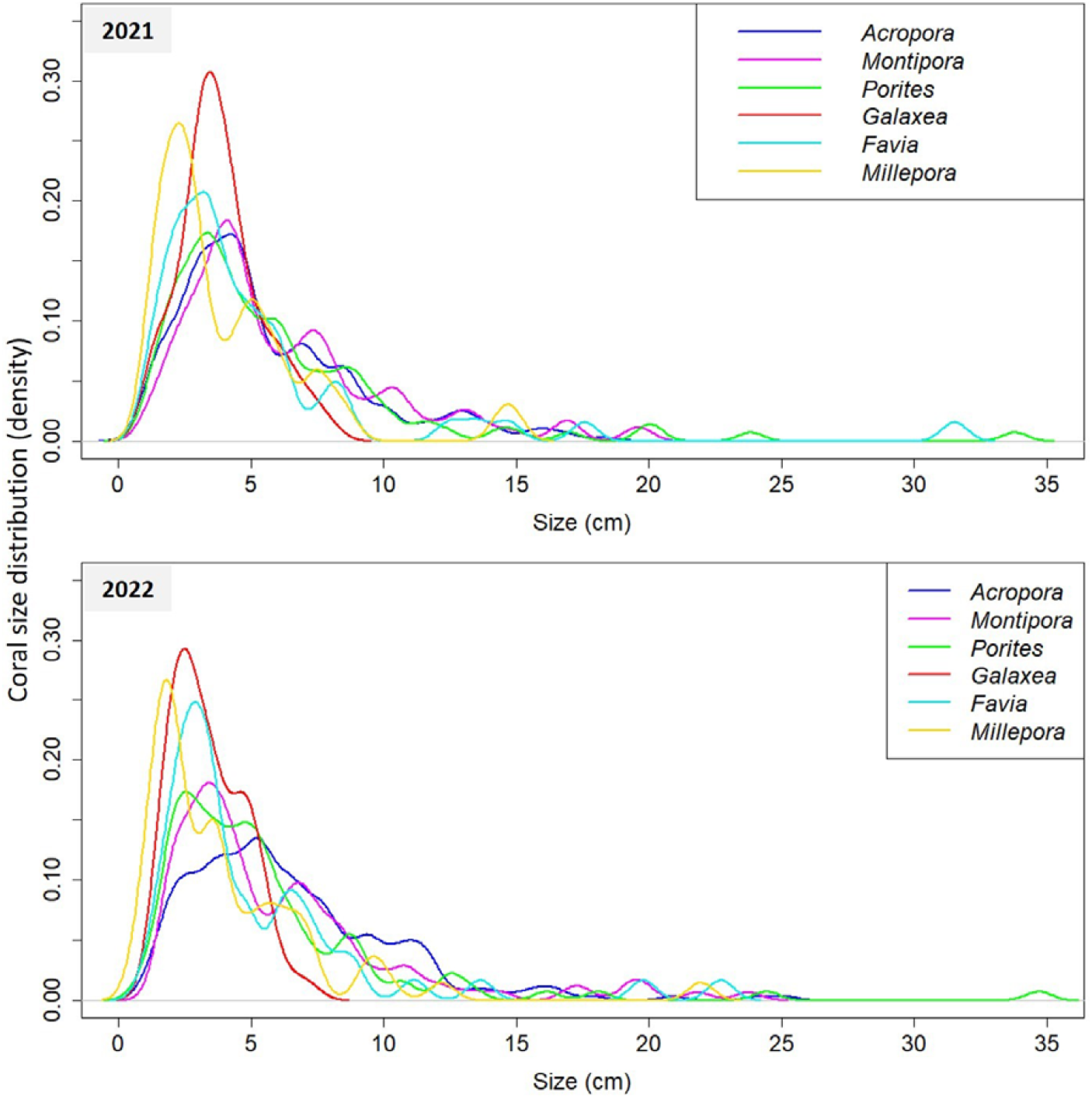
Size distributions of populations of the six dominant coral genera in 2021 and 2022. See S5 Figure for inter-annual differences in size-distributions.

### Coral demographic performances

Coral survival was lowest at small sizes and increased with colony size, except for *Favia* in which the probability of survival was consistently >90% across the size-range (Fig. 5A). The other coral genera showed different degrees of size-refuging, i.e. when survival increases with size as seen in many species (Madin et al. 2014; Kayal et al. 2015; Ellner et al. 2016), with *Porites* showing the highest survival, >90% for colony sizes above 5 cm and ∼98% beyond 10 cm. In contrast, survival was lowest for *Millepora* with a 70% chance of survival for a colony size of 10 cm. *Galaxea* had the lowest survival rate at small sizes, reaching the 50% survival size threshold at a mean colony diameter of 2.5 cm. Comparatively, *Acropora* and *Porites* colonies achieved a 50% survival probability at a size of ∼1.5 cm, and the other genera at even smaller sizes.

**Fig. 5.**
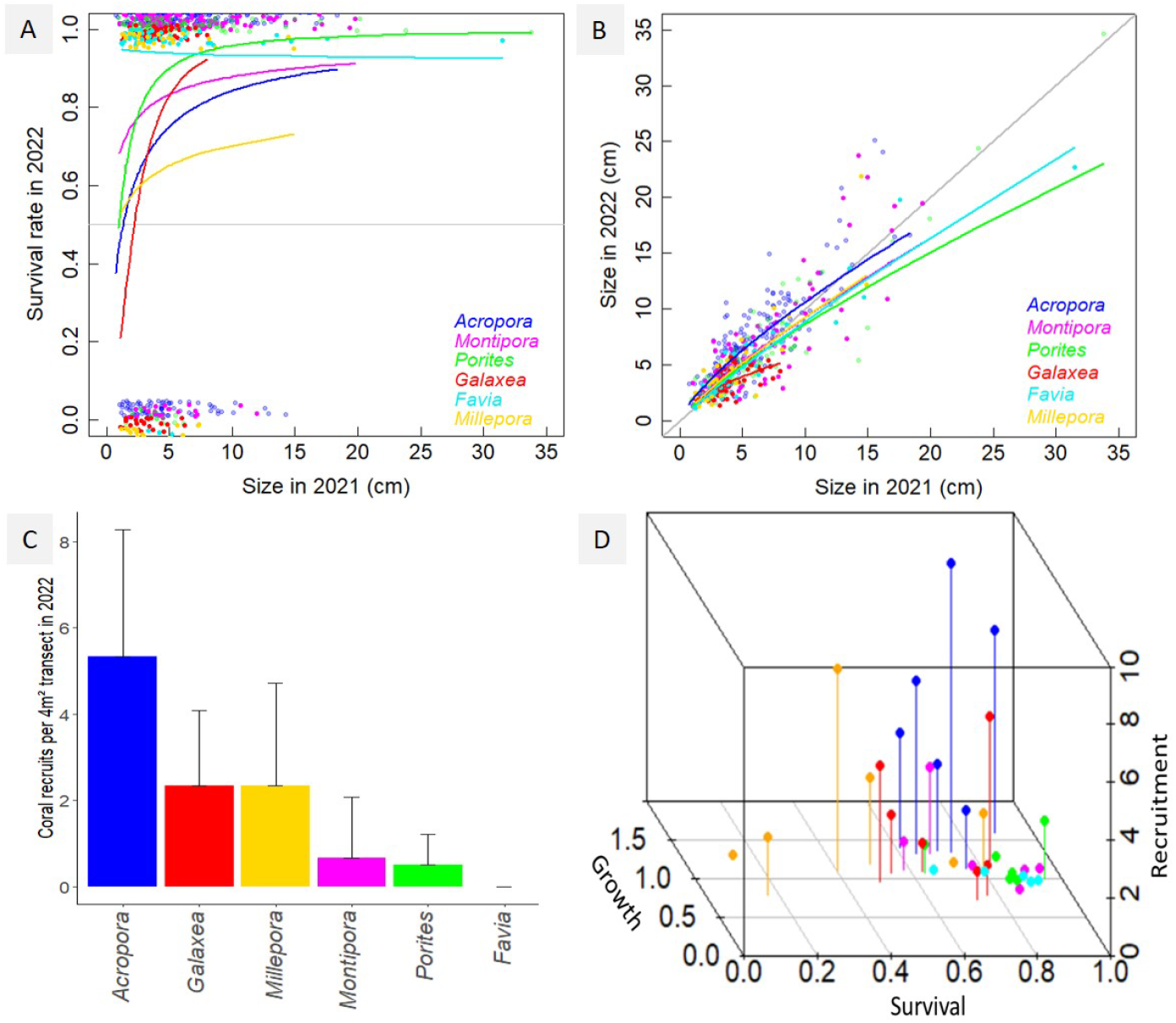
Individual coral performances in survival (A), growth (B), and recruitment (C) for the six dominant genera, and combined (D). Survival and growth rates are expressed as a function of colony sizes (mean coral diameter), with points representing individual observations, and curves estimated patterns from Generalized Linear Mixed-effect Models. Gray line in survival plot (A) indicates the 50% survival threshold. Note that data points in (A) were jittered around 1 (i.e. survival) and 0 (i.e. mortality) to ease visual distinction of individual observation points. Grey line in growth plot (B) indicates the identity threshold (Size_2022_=Size_2021_), values above the line indicate positive growth, and below coral shrinkage. Recruit abundances are expressed as mean ± standard deviation. The three-dimensional plot (D) combines average coral genera survival, growth, and recruitment performances information per transect into a three-dimensional space for a visualization of differences in coral life history. Survival, growth, and recruitment rates in (D) are in the same units as in (A), (B), and (C).

In all genera, coral relative growth was highest at small sizes and shifted to colony shrinkage at larger sizes, though at rates among taxa (Fig. 5B). *Acropora* had the highest growth at all sizes, with colony shrinkage becoming predominant beyond an average size of 13 cm. In contrast, *Galaxea* had the lowest growth rate with many cases of colony shrinkage, even at small, mean diameter <5 cm stages. The other genera showed intermediate patterns with, on average, positive growth until a size of ∼7.5 cm, beyond which they tended to shrink.

For recruitment, 11 ±4 SD coral recruits per 4 m^2^ transect were recorded, the majority being *Acropora* (5 ±3 SD; Fig. 5C). No recruits were recorded for *Favia*, while the other genera showed intermediate levels with 1-2 recruits per transect.

Summarizing coral demographic performances in survival, growth, and recruitment in a three-dimensional space enables a visual distinction of species’ life history characteristics (Fig. 5D). High recruitment and growth rates mark *Acropora. Montipora, Porites*, and *Favia* produce low numbers of recruits that have higher chances of survival, whereas the calcifying hydrozoan *Millepora* (a.k.a. fire-coral), compensates low survival with intermediate recruitment rates. Galaxea shows intermediate features.

## Discussion

### Characterizing coral demographic performances

Demographic studies are foundational to comprehend drivers of species dynamics and evaluate community responses to changing environments (Ellner et al. 2016). Yet, limitations in data acquisitions have constrained their large-scale applications in population ecology and ecosystem conservation. This is the case in New Caledonia where lies one of the world’s largest and most prolific coral reef systems (Andréfouët et al. 2009), but where no studies on coral demographic performances existed. At a time when environmental changes and impacts to ecosystems press for increased quantitative knowledge on species dynamics and their drivers, our relatively simple digital approach enables extracting data on community abundance and composition and the dynamics of individuals therein from video transects, facilitating resource-effective, image-based demographic investigations. Applied to our underwater, outer-reef coral community in New Caledonia, we characterize community composition and demographic performances of the dominant reef-building coral taxa over a year, providing insight into species life histories and constraints to their demographics.

The studied coral community comprised 26 genera occupying the reef substrate with a density of 36 colonies per m^2^, with six dominant genera representing >75% of corals, by order of abundance: *Acropora, Montipora, Porites, Galaxea, Favia*, and *Millepora*. Coral populations were predominantly constituted of small, <5 cm colonies, with *Acropora, Montipora*, and *Porites* showing higher proportions of larger corals. These six genera typically dominate coral communities in Southwest Pacific reefs, as well as the global reefscape (Adjeroud et al. 2018, 2019; Darling et al. 2019; Dietzel et al. 2021).

Taxonomic differences in coral abundance, size, survival, growth, and recruitment reveal a variety of life histories among dominant genera and shed light on the diverse evolutionary pathways towards the ecological success of dominant coral taxa – a segregation of ecological niches promoting species coexistence and high levels of biodiversity (Tokeshi 2009; Holt 2017; Kayal & Adjeroud 2022). The most abundant genus, *Acropora* (representing 30% of surveyed corals) prevailed in recruitment and growth, which is unsurprising as these are two prominent features of this taxon reported in various regions (Kayal et al. 2015; Hughes et al. 2019; Adjeroud et al. 2022). In contrast, higher survival compensated for lower recruitment and growth in *Montipora, Porites*, and *Favia*. Massive growth forms of *Porites* and *Favia* are archetypes of the stress-tolerant life-strategy in corals, typically dominating reefs in extreme and altered environments, whereas *Montipora* often exhibits intermediate life history features between *Acropora* and *Porites*, although within genera differences also prevail (Kayal et al. 2018; Darling et al. 2019). *Galaxea* and *Millepora* also differentiated from the other taxa, achieving abundant populations despite relatively low survival and intermediate recruitment rates.

Many corals exhibited colony shrinkage rather than growth. As other coral reefs in the South Pacific, our study site on the outer reef slope is subject every year to several events of strong south-west storm swells breaking apart coral colonies and occasionally chunks of the reef substrate (Fig. 6). While recurrent hydrodynamic stress appears clearly as a limiting factor for coral development on these outer reef slope sites, further investigation remains necessary to evaluate the degree to which intrinsic species life history traits, and extrinsic abiotic and biotic environmental conditions in concert, drive coral community dynamics. Prior studies have highlighted taxonomic differences in coral vulnerability to physical dislodgement and fragmentation (Madin et al. 2014; Kayal et al. 2015), with some taxa using fragmentation as a strategy for asexual propagation. This is the case for *Acropora, Galaxea*, and *Millepora*, the three genera showing the highest recruitment rates on our study site recurrently impacted by strong waves. Characterized by digitated growth forms (Fig. 6), *Acropora* branches break relatively easily into loose fragments that show a high capacity to survive and reattach to the substrate (Kayal et al. 2015). Similarly, *Galaxea* colonies often split into detached individual polyps (Fig. 7), and *Millepora* pieces into free branches and columns that are dispersed by waves (Dubé et al. 2021). Nevertheless, recent investigations in our study system indicate that the outer reef receives lower rates of larval settlement than nearby lagoon sites, and that several coral taxa exhibit lower competitive performances there as compared to other reef environments (Adjeroud et al. 2022; Kayal & Adjeroud 2022). Expanding coral demographic surveys in time and space, and complementing statistical analyses with simulation-based modelling approaches will help identify key processes controlling coral demographic success, and estimate critical stress thresholds for coral resilience (Madin et al., 2014; Kayal et al. 2018; Riegl et al. 2018; Carlot et al. 2021). By improving fieldwork time-efficiency and thus expansion of population and community dynamics studies, our image-based approach to characterizing species demographic performances will facilitate this endeavor.

**Fig. 6.**
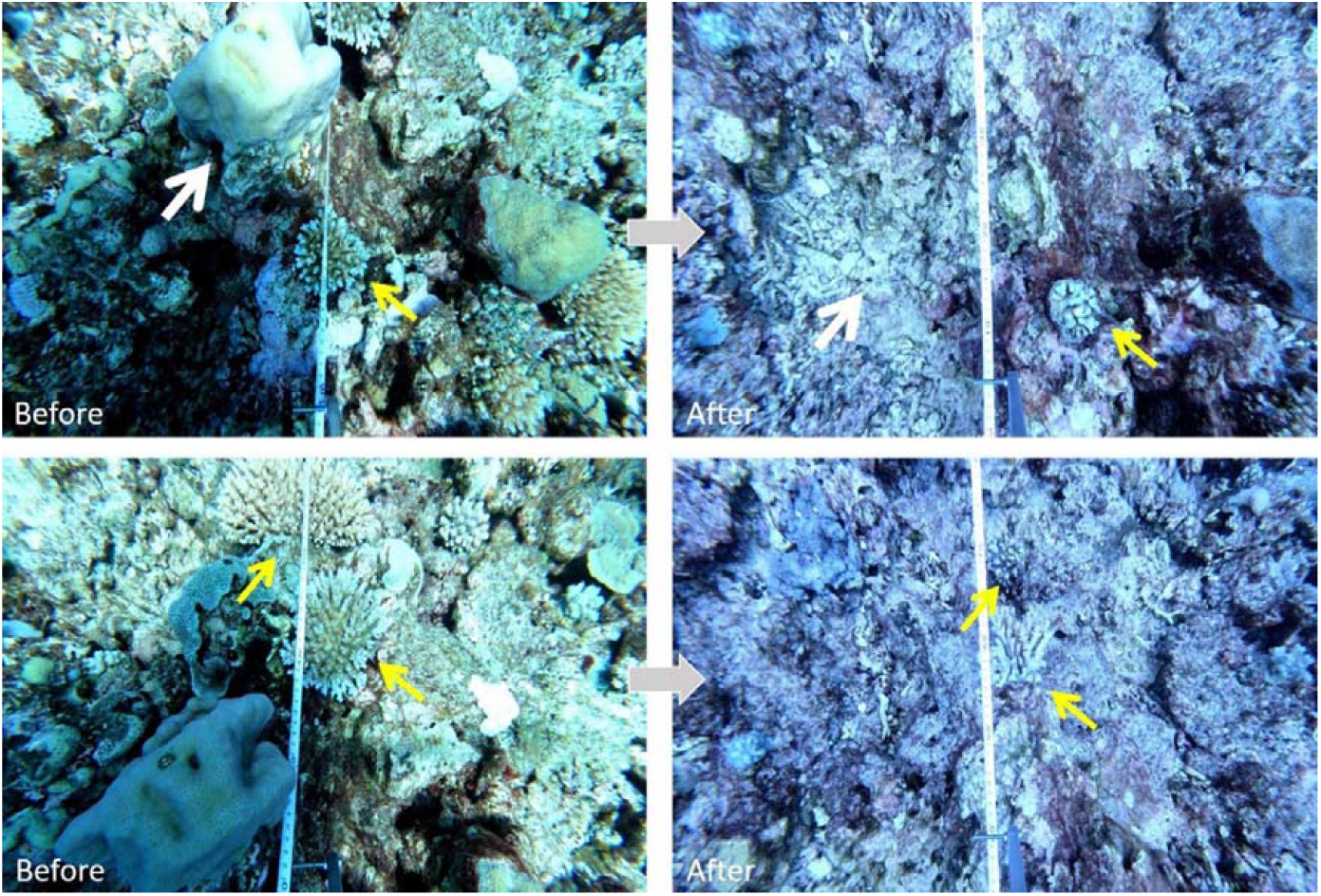
Photographs showing two portions of the reef impacted by strong waves. Yellow arrows indicate branching *Acropora* colonies shrinking in size due to fragmentation (i.e. broken branch tips following high hydrodynamic stress). White arrows indicate the position of a massive *Favia* colony that was blasted away along with a chunk of the reef substrate.

**Fig. 7.**
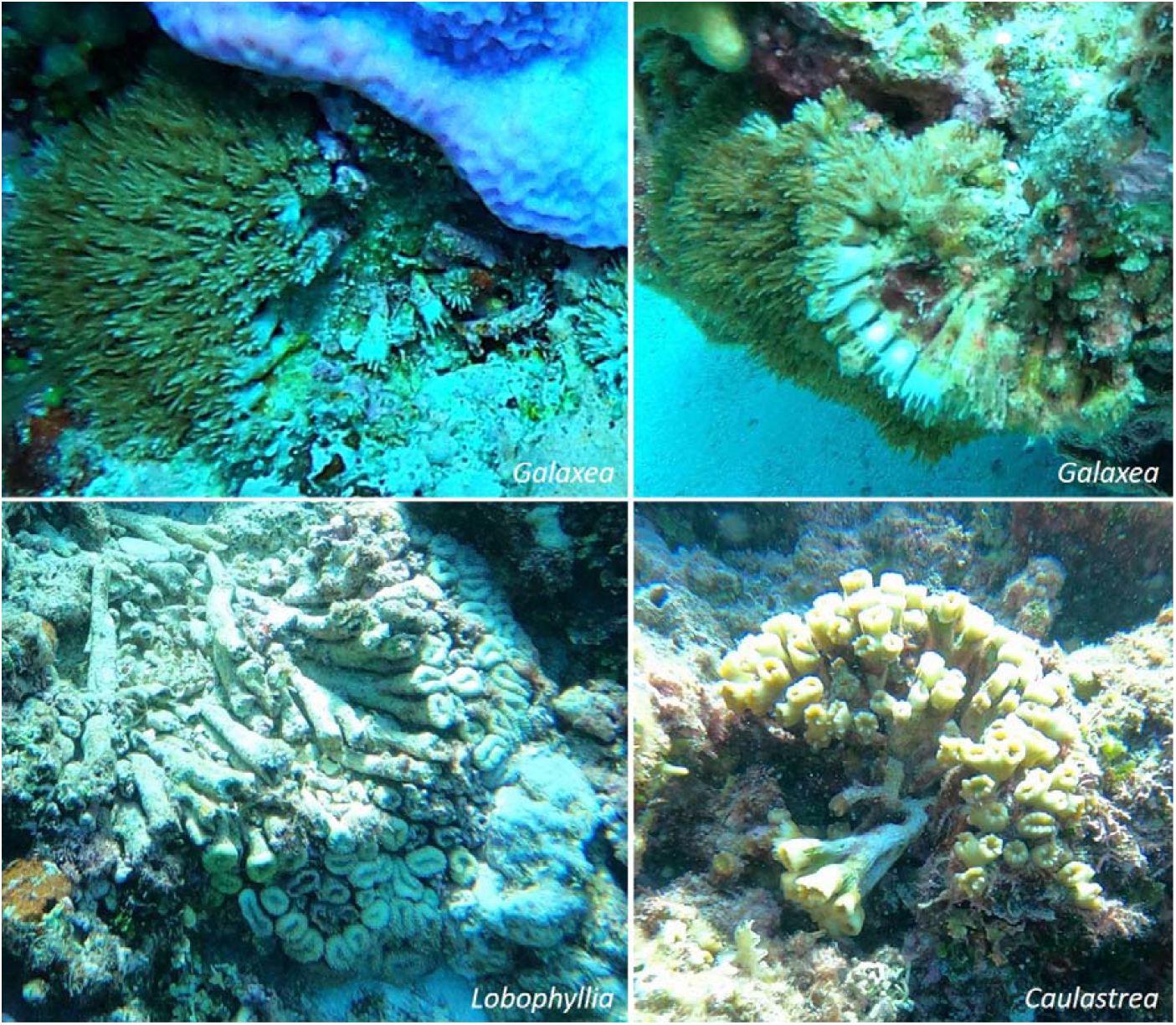
Photographs showing asexual propagation of loose polyps by massive *Galaxea* and other coral genera.

### Accelerating demographic investigation using imagery

Because data acquisition during field surveys is often restricted by time constraints, imagery based approaches can greatly optimize data collection and archiving, particularly for extreme and remote environments where immersion is limited. Our digital approach helps address the growing need of managers for accessible ecological diagnostics, including studies that track species’ individual performances in key demographic processes such as survival, growth, reproduction, and migration to characterize demographic bottlenecks and comprehend community resilience (Ellner et al. 2016; Kayal et al. 2018; Riegl et al. 2018). By operationalizing a transition of data acquisition tasks from in situ note taking to imagery-based annotations, our approach complements ongoing technological developments that rely on the digitalization of the natural world such as photogrammetry (Urbina-Barreto et al. 2021), and automatization of data acquisitions and treatments such as artificial intelligence (González-Rivero et al. 2020). We particularly advocate for applications in the preservation of declining key ecological communities threatened by near-future collapse, such as reef-building corals (Dietzel et al. 2020; Jamil et al. 2021).

## Acknowledgments

This study is part of the Track Changes project, which was supported by the French Laboratory of Excellency CORAIL through a starting grant and a graduate student stipend, and the French Ministry of Foreign Affairs through the regional cooperation grant Fonds Pacifique. We thank Yassine Majoul for help with computer coding, and staff from the Institute of Research for Development for assistance for boating and diving.

## Supplementary material captions

**S1 Figure** Location of our study site at an outer reef slope in New Caledonia.

**S2 Appendix** Python code for mapping and measuring corals in video transects.

**S3 Table** Population abundances (n per 4 m^2^) and relative genera contributions (%) in our surveys of coral communities in 2021 and 2022.

**S4 Table** Kolmogorov-Smirnov test results comparing size distributions of the six dominant coral genera in 2021 and 2022.

**S5 Figure** Size distributions of populations of the six dominant coral genera in 2021 and 2022.

## Notes

### Competing Interest Statement

The authors have declared no competing interest.

### Summary of Updates

Following off-topic comments by anonymous reviewers, minor changes were performed to the Abstract to clarify the approach: data extractions from video-transects are performed manually as a digital version of what is traditionally performed in situ, though the approach can be automatized in the future via association with appropriate algorithms. Similarly, additional details were added to the data analysis section. Two figures were moved from supplementary material to the core of the manuscript. Few additional references were also added. Figures are now displayed within the text for a more pleasant read. Enjoy!

