## Supplementary material for "Coral demographic performances in New Caledonia, a video transect approach to operationalize imagery-based investigation of population and community dynamics"

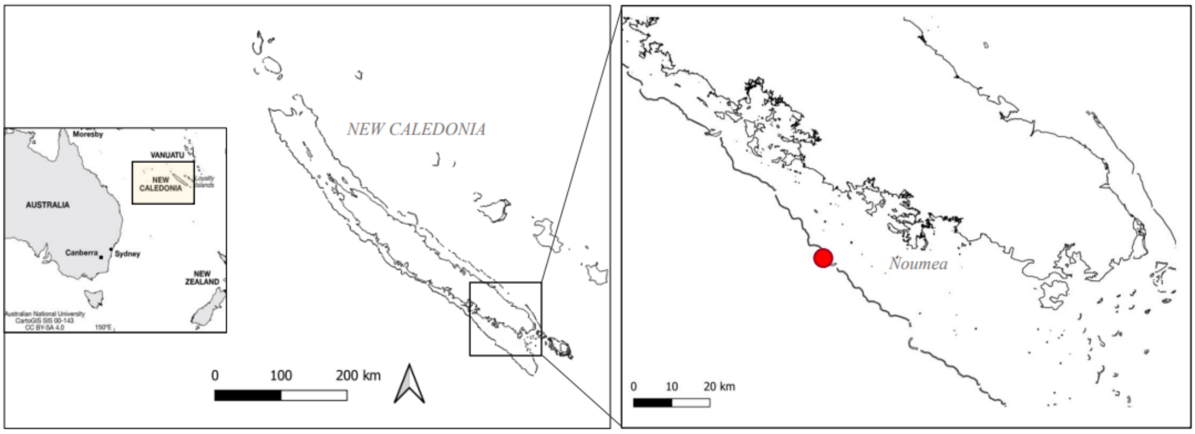


**S1 Figure** Location of our study site at an outer reef slope in New Caledonia.

**S2 Appendix** Python code for mapping and measuring corals in video transects.

```
import math
from optparse import Option
from tkinter import *
from cmath import pi
from colorsys import yiq_to_rgb
from pickle import FALSE
from re import X
import cv2
import math
import os
import tkinter
import tkinter.simpledialog
from tkinter import *
import statistics
import numpy as np
from pynput import mouse
import pandas as pd
import keyboard
import csv


## colors ##
colors = {'blue': (255, 0, 0), 'green': (0, 255, 0), 'red': (0, 0, 255),
'yellow': (0, 255, 255),'magenta': (255, 0, 255),
'cyan': (255, 255, 0), 'white': (255, 255, 255), 'black': (0, 0, 0),
'gray': (125, 125, 125)}

def emptystring(string):
    answer = False
    if len(string) == 0:
        answer = True
    elif not string:
        answer = True
    elif string.isspace():
        answer = True
    return answer

def mainMenu():
    videosFolder = os.listdir(os.getcwd() + "/videos")

    def selected():
        chosenVid = clicked.get()
        videoPlayer(chosenVid)

    window=Tk()
    window.title('MeasureVids')
    window.geometry("470x270") #"960x540"

    clicked = StringVar()
    clicked.set(videosFolder[0])
    drop = OptionMenu(window, clicked, *videosFolder)
    drop.pack(pady=20)

    selection = Button(window, text="Choose this vid!", command=selected)
    selection.pack()
```

```
    window.mainloop()

def videoPlayer(filename):
    pathCSV = os.getcwd()+"/CSVs/"
    path = os.getcwd()+"/videos/"+filename #before: create a folder named "videos"
    pathscreenshots = os.getcwd()+"/screenshots/"+filename+'/'
    if not os.path.exists(pathscreenshots):
        os.makedirs(pathscreenshots) #create folder Screenshots
        print('Creating a screenshot folder for ' + filename)
    if not os.path.exists(pathCSV): #create folder CSV
        os.makedirs(pathCSV)
        print('Creating a CSV folder for ' + filename)

    cap = cv2.VideoCapture(path)
    fps = cap.get(cv2.CAP_PROP_FPS) #frame per second

    ## global variables ##

    image_list = []
    coral_id_list = []
    genus_list = []
    morpho_list = []
    length_list = []
    width_list = []
    midlength_list = []
    midwidth_list = []
    centroid_list = []
    frame_list = []
    time_list = []
    scale_list = []
    col_tr_px_list = []
    col_tr_cm_list =[]
    position_x_list = []
    angle_rota_list = []
    dynamic_list = []
    coral_id_long_list = []
    Dm_list = []
    pointA_list = []
    pointB_list = []
    pointC_list = []
    pointD_list = []
    penta_list = []
    fusion_fission_list = []
    size_difference_list = []
    frameNb = 0
    count=0
    coord = []
    roiPts = []
    scalepts = []
    choose_smp_ef = 40 #sampling effort over 0.8 meter width (40 centimeters on the left and on the right)

    # opening the CSV file
    # Comparison of csv transects in 2021 and 2022
    csvFile = pd.read_csv('Transect_6_2021.csv', sep=';')
    coral_id_2021 = csvFile['Individual_ID'].values.tolist()
    individual_2021 = csvFile['Individual_ID_short'].values.tolist()
```

```
size_2021 = csvFile['Mean_diameter'].values.tolist()
coral_morpho_id = csvFile['Morpho'].values.tolist()
coral_genus_id = csvFile['Genus'].values.tolist()


#program to scale each image relative to the transect tape

def scaling(event,x,y,flags,params):
    global imagetmp
    imagetmp = image
    global pixelScale
    global xAB, yAB

    if event == cv2.EVENT_RBUTTONDOWN: #if right click
        scalepts.append((x, y)) #add coordinates
        #print("scale pts:", x, y)
        #print(scalepts)
        cv2.circle(imagetmp, (x, y), 2, (255, 0, 0), -1) #display a circle

        if len(scalepts) == 2: #when two points have been picked out
            cv2.line(imagetmp, scalepts[0], (x, y), (255, 0, 0), 2) #line between the first and the second point
            xAB = scalepts[0][0] #x of the first point
            yAB = scalepts[0][1] #y of the first point
            xBB = scalepts[1][0] #x of the second point
            yBB = scalepts[1][1] #y of the second point
            pixelDimension = math.sqrt((xBB-xAB)**2 + (yBB-yAB)**2) #nb of pixels of the scale
            #print("ok", pixelDimension)
            metricDimension = tkinter.simpledialog.askfloat(title="Calibrate", prompt="Scale (cm):") #ask the
equivalent in centimeters
            pixelScale = metricDimension/pixelDimension #cm/pixel
            print("Size in cm/px :" + str(pixelScale))

            #if you want to add a horizontal line to restrain the width of the frame considered
            ymoy = int((yAB+yBB)/2) #at the middle of the scale
            xmoy = int((xAB+xBB)/2) #at the middle of the scale
            smp_ef = int(choose_smp_ef/pixelScale) #choose_smp has to be precised (in centimeters) at the
beginning
            cv2.line(imagetmp, (xmoy,ymoy), (xmoy+smp_ef, ymoy), (50, 0, 155), 2)
            cv2.line(imagetmp, (xmoy,ymoy), (xmoy-smp_ef, ymoy), (50, 0, 155), 2)

            scalepts.clear()

            return pixelScale


#program to annotate the frame

def annot(event, x, y, flags, params):
    global imagetmp
    imagetmp = image
    global xA, yA, xB, yB
    global Xcentroid1, Ycentroid1, lengthpx, length

    global count_unknown
    count_unknown = 0 #to count the unknown individuals

    if event == cv2.EVENT_LBUTTONDOWN: #if left click
```

```
roiPts.append((x, y)) #add coordinates
#print("roi just added:", x, y)
#print("roi points=", roiPts)
cv2.circle(imagetmp, (x, y), 2, (0, 0, 200), -1) #colored dots
if len(roiPts) == 2: #if two points have been picked out (ie length)
    cv2.line(imagetmp, roiPts[0], (x, y), (255, 0, 255), 2)
    #magenta line between the first and the second point
    xA = roiPts[0][0]
    yA = roiPts[0][1]
    xB = roiPts[1][0]
    yB = roiPts[1][1]
    lengthpx= math.sqrt((xB-xA)**2 + (yB-yA)**2)
    #print("length in px is:", lengthpx)
    length = pixelScale * lengthpx #length in cm
    print("real length in cm is:", length)
    length_list.append(round(length,1)) #round to one number after comma

    Xcentroid1 = int((xA+xB)/2) #midlength
    Ycentroid1 = int((yA+yB)/2) #midlength

    midlength_list.append((Xcentroid1,Ycentroid1))
    cv2.circle(imagetmp, (Xcentroid1, Ycentroid1), 2, (0, 255, 255), -1) #circle on the midlength

if len(roiPts) == 4: #4 points
    cv2.line(imagetmp, roiPts[2], (x, y), (50, 255, 50), 2)
    #line between the third and the fourth point
    xC = roiPts[2][0]
    yC = roiPts[2][1]
    xD = roiPts[3][0]
    yD = roiPts[3][1]
    widthpx = math.sqrt((xD-xC)**2 + (yD-yC)**2)
    #print("width in px is:", widthpx)
    width = pixelScale * widthpx
    print("real width in cm is:", width)
    width_list.append(round(width,1))

    Xcentroid2 = int((xC+xD)/2) #midwidth
    Ycentroid2 = int((yC+yD)/2) #midwidth

    pointA_list.append((xA,yA))
    pointB_list.append((xB,yB))
    pointC_list.append((xC,yC))
    pointD_list.append((xD,yD))

    midwidth_list.append((Xcentroid2,Ycentroid2))
    #cv2.circle(imagetmp, (Xcentroid2, Ycentroid2), 2, (0, 255, 255), -1) #circle on the midwidth

    X_centroid = int((Xcentroid1+Xcentroid2)/2) #barycentre - could be considered as the centroid?
    Y_centroid = int((Ycentroid1+Ycentroid2)/2) #barycentre - could be considered as the centroid?
    centroid_list.append((X_centroid, Y_centroid))
    #cv2.circle(imagetmp, (X_centroid, Y_centroid), 3, (0, 0, 255), -1) #midlength

    #ellipse drawing
    length_int = int(lengthpx/2) #divided by two otherwise too large
    width_int = int(widthpx/2)
    axesLength = (length_int, width_int)
    if xB == xA:
```

```
    xB = xB+1 #cannot divide by zero
fkslope = (yB-yA)/(xB-xA)
if fkslope < 0: #properties of arctan: if arg is negative then results is underestimated
    angle = 180 - ((np.arctan(math.sqrt((yA-yB)**2)/math.sqrt((xA-xB)**2)))*(180/pi))
else:
    angle = (np.arctan(math.sqrt((yA-yB)**2)/math.sqrt((xA-xB)**2)))*(180/pi)
#print(angle)
angle_rota_list.append(angle)
cv2.ellipse(imagetmp, (Xcentroid1,Ycentroid1), axesLength, angle, 0, 360, (0, 100, 205), 2) #centroid =
middle of the length

Dm_cm = (length+width)/2 #mean diameter
Dm_list.append(Dm_cm)
roiPts.clear()

identification = tkinter.simpledialog.askstring(title="Identification", prompt="'genus_morpho':")
while emptystring(identification):
    identification = tkinter.simpledialog.askstring(title="Identification", prompt="'genus_morpho':")
font = cv2.FONT_HERSHEY_SIMPLEX

word_split = identification.split('_') #split genus and morpho
genus = word_split[0]
morpho = word_split[1]
if len(word_split) == 3: #if it is 'genus_morpho_comment' (3 components)
    fu_fi = word_split[2] #fusion or fission
else:
    fu_fi = 'NA'

genus_list.append(genus)
if genus == "unk": #if unknown
    count_unknown +=1 #to know how many unknown are identified; can be #
#ID_list.append(str(uuid.uuid4())[:8]) #random coral ID
frame_list.append(frameNb) #add frame n°
image_list.append(filename+'_'+idCapture+".png") #add image name
time_list.append(time) #add time in the video
scale_list.append(pixelScale) #add scale in cm/px

##position Y (distance between the colony centroid and the transect) --> relative to the transect width
distance_col_tr_px = math.sqrt((xAB-Xcentroid1)**2)
#print("distance from the colony to the transect in px is:", distance_col_tr_px)
col_tr_px_list.append(round(distance_col_tr_px))

penta_list.append((xAB,yAB)) #coordinates of the transect tape

distance_col_tr_cm = distance_col_tr_px * pixelScale
print("Real distance from the colony to the transect in cm is:", distance_col_tr_cm)
if Xcentroid1 < xAB: #colony on the left side of the transect
    distance_col_tr_cm = - distance_col_tr_cm #negative on the left

col_tr_cm_list.append(round(distance_col_tr_cm))

##position X (position on the transect (100cm, 120cm,...)) --> relative to the transect length
position_tr = tkinter.simpledialog.askfloat(title="Transect graduation", prompt="Position on the
transect (cm):")
#print("Coral colony located at:", position_tr)
position_x_list.append(position_tr)
coral_id = genus+'_'+morpho+'_'+str(round(position_tr))
```

```
            if morpho == "ma":
                morpho = 'massive'
            elif morpho == "sm":
                morpho = 'submassive'
            elif morpho == 'fo':
                morpho = 'foliose'
            elif morpho == 'br':
                morpho = 'branched'
            elif morpho == 'en':
                morpho = 'encrusting'
            elif morpho == 'ta':
                morpho = 'tabular'
            elif morpho == 'so':
                morpho = 'solitary'
            print("morpho is:", morpho)
            morpho_list.append(morpho)

            coral_id_list.append(coral_id) #ID short is: genus_morpho_x
            fusion_fission = 'NA'
            size_difference = 'NA'

            # coral_id_long = coral_id+'_2021'
            # coral_id_long_list.append(coral_id_long)
            #dynamic = 'NA'
            #dynamic_list.append(dynamic)

            #comparison to 2021
            if coral_id in individual_2021: #if the coral has been seen before (coral of 2022 % list 2021)
                coral_id_long = coral_id+'_2021'
                coral_id_long_list.append(coral_id_long)

                for i in range(0, len(coral_id_2021)):
                    name = coral_id_2021[i]
                    if coral_id_long == name: #if they have the same name (on row i)
                        if Dm_cm >= size_2021[i]: #if coral 2022 is larger than 2021
                            if 'fu' in fu_fi:
                                dynamic = 'fusion_growth'
                                dynamic_list.append(dynamic)
                                fusion_fission = tkinter.simpledialog.askstring(title="Fusion", prompt="Fusion with which
colony? 'genus_morpho_x_year'")
                                print("the coral has grown and merged with", fusion_fission)
                            elif 'fi' in fu_fi:
                                dynamic = 'fission_growth'
                                dynamic_list.append(dynamic)
                                fusion_fission = tkinter.simpledialog.askstring(title="Fission", prompt="Fission from which
colony? 'genus_morpho_x_year'")
                                print("the coral has grown and split into", fusion_fission) #takes the larger coral for fi
                            else:
                                dynamic = "growth"
                                dynamic_list.append(dynamic)
                                print("the coral has grown")

                        else: #if the coral has shrunk
                            if 'fu' in fu_fi:
                                dynamic = 'fusion_shrinkage'
                                dynamic_list.append(dynamic)
```

```
                        fusion_fission = tkinter.simpledialog.askstring(title="Fusion", prompt="Fusion with which
colony? 'genus_morpho_x_year'")
                        print("the coral has shrunk and merged with", fusion_fission) #the smaller colony
                    elif 'fi' in fu_fi:
                        dynamic = 'fission_shrinkage'
                        dynamic_list.append(dynamic)
                        fusion_fission = tkinter.simpledialog.askstring(title="Fission", prompt="Fission from which
colony? 'genus_morpho_x_year'")
                        print("the coral has shrunk and split into", fusion_fission)
                    else:
                        dynamic = "shrinkage"
                        dynamic_list.append(dynamic)
                        print("the coral has shrunk")
                size_difference = Dm_cm - size_2021[i]

                print('size difference between 2021 and 2022:', size_difference)



        else: #coral not seen in 2021; new
            coral_id_long = coral_id+'_2022' #new id with 2022
            coral_id_long_list.append(coral_id_long)

            dynamic = tkinter.simpledialog.askstring(title="Dynamic", prompt="Coral status: (r: recruit, mi:
missed, fu: fusion, fi: fission, mig : migrant)")
            if dynamic == 'r':
                dynamic = 'recruit'
                print("This coral is a new recruit") #small colony
            elif dynamic == 'mi':
                dynamic = 'missed'
                print("This coral was missed") #camera angle, resolution, edge, etc.
            elif dynamic == 'fi':
                dynamic = 'fission'
                fusion_fission = tkinter.simpledialog.askstring(title="Fission", prompt="Fission from which
colony? 'genus_morpho_x_year'")

                print("This coral comes from a fission from the colony:", fusion_fission) #the smaller part is
considered
            elif dynamic == 'mig':
                dynamic = 'migrant'
                print("This coral is a new fragment or a migrant") #for solitary corals (ex: fungia), fragments...
            elif dynamic == 'fu':
                dynamic = 'fusion'
                fusion_fission = tkinter.simpledialog.askstring(title="Fusion", prompt="Fusion with which colony?
'genus_morpho_x_year'")

                print("This coral comes from a fusion from the colony:", fusion_fission)
            dynamic_list.append(dynamic)
        fusion_fission_list.append(fusion_fission)
        size_difference_list.append(size_difference)

        # screen_width = image.shape[1]
        # text_x_pos = None
        # text_y_pos = y
        # if x < (screen_width/2): #if you annotate a colony on the left:
        #     text_x_pos = int(x + 5) #text on the right
        # else:
        #     text_x_pos = int(x - 5) #annotation on the right --> text on the left
```

```
            cv2.putText(imagetmp, coral_id_long, (xA,yA), font, 0.5, (0, 255, 255), 2) #coral_id_year next year

        print("You may select your next point!")

    def main(event,x,y,flags,params): #the main program used

        scaling(event,x,y,flags,params) #call for scaling first
        annot(event,x,y,flags,params) #call for annotationg function

    while cap.isOpened(): #while the video is running

        ret, frame = cap.read()
        resized = cv2.resize(frame, (960,540), interpolation = cv2.INTER_LINEAR) #small screen: 960,540 // bigger
screen: 1366,768
        video = cv2.imshow("Video LIT test", resized) #show video resized

        if ret == True:  #if the video is played:
            key = cv2.waitKey(1) #1000/fps)) #1000ms/ nb frame per s
            frameNb +=1

            if key == ord('e'): #if press E

                #frameNb = cap.get(CAP_PROP_POS_FRAMES)
                idCapture = str(frameNb) #str(uuid.uuid4())[:8] #uuid gives a random ID (chr)
                time = frameNb/fps #time of the frame
                print("Frame number:" + idCapture)
                print("Time (s):" + str(time))
                imgname = pathscreenshots+filename+'nonannot_'+idCapture+".png"

                cv2.imwrite(imgname, frame) #save screen
                imagefirst = cv2.imread(imgname)
                image = cv2.resize(imagefirst, (1100,575), interpolation = cv2.INTER_LINEAR) #resized
                print("Please press left to get the coordinates, press right to calibrate")

                cv2.namedWindow('Image mouse')
                cv2.imshow('Image mouse', image)
                cv2.setMouseCallback('Image mouse', scaling)

                while True:
                    # Show image 'Image mouse':
                    cv2.namedWindow('Image mouse')
                    cv2.imshow('Image mouse', image)  #shows the frame considered
                    cv2.setMouseCallback('Image mouse', main)

                    if cv2.waitKey(int(1000/fps)) & 0xFF == ord('q'): #press q to quit
                        coord.clear()
                        imgnameAnnot = pathscreenshots+filename+'annot_'+idCapture+".png"
                        cv2.imwrite(imgnameAnnot, imagetmp) #save screen annoted

                            # Create a dictionary using lists
                        data = {'Individual_ID' : coral_id_long_list, 'Individual_ID_short' : coral_id_list, 'Genus':genus_list,
'Morpho':morpho_list, 'Length_cm ': length_list, 'Width_cm': width_list, 'Mean_diameter': Dm_list,
'Position_Y_cm': col_tr_cm_list, 'Position_X_cm': position_x_list, 'Dynamic': dynamic_list, 'Size_difference (cm)'
: size_difference_list, 'Fusion/Fission' : fusion_fission_list,'Position_Y_px': col_tr_px_list,'Image': image_list,
'Frame_Number': frame_list, 'Frame_Time': time_list, 'Scale_cm/px' : scale_list, 'Centroid' : centroid_list,
'Mid_Length' : midlength_list, 'Mid_Width' : midwidth_list, 'Ellipse_Angle' : angle_rota_list, 'Point_A':
pointA_list,'Point_B': pointB_list,'Point_C': pointC_list,'Point_D': pointD_list, 'Scale_position' : penta_list}
```

```
            # Create the Pandas DataFrame
            df = pd.DataFrame(data)
            #print()
            #print(df)
            #print()
            # Export the dataframe to a csv file
            df.to_csv(pathCSV +filename+'.csv', index = False, header=True, sep=';') #use ';' as sep even
though ',' is the real separator
            #https://stackoverflow.com/questions/63244161/why-is-pandas-to-csv-comma-seperating-not-
working

            if count_unknown != 0:
              print('You have',count_unknown, 'coral unknown on the frame', frameNb)
            break

        # Destroy all generated windows:
        cv2.destroyAllWindows()
        count += 1

    #### --------------------  at the end of the video ---------------------------- ####

      if key == 27: #press esc to quit
        print("End of the video")
        break

    else:
      break

  cap.release()
  cv2.destroyAllWindows()


  ## uncomment when you want to work on the second year
  # below

  #at the end of the program, if a coral in 2021 has not been encountered in 2022:

  v = len(coral_id_2021) #list of ID of the transect 6 2022
  df_f = df
  for q in range(v):
    if coral_id_2021[q] not in coral_id_long_list: #check on genus_morpho_x_year
      new_row = {'Individual_ID' : coral_id_2021[q], 'Individual_ID_short': individual_2021[q],
'Genus':coral_genus_id[q], 'Morpho':coral_morpho_id[q], 'Length_cm ': 0, 'Width_cm': 0, 'Mean_diameter': 0,
'Position_Y_cm': 'NA', 'Position_X_cm': 'NA', 'Dynamic' :'Absent', 'Size_difference (cm)' : 'NA', 'Fusion/Fission' :
'NA', 'Position_Y_px': 'NA','Image': 'NA', 'Frame_Number': 'NA', 'Frame_Time': 'NA', 'Scale_cm/px' :
'NA','Centroid' : 'NA', 'Mid_Length' : 'NA', 'Mid_Width' : 'NA', 'Ellipse_Angle' : 'NA', 'Point_A': 'NA','Point_B':
'NA','Point_C': 'NA','Point_D':'NA', 'Scale_position': 'NA'}
      df_new_row = pd.DataFrame(new_row, index=[0])
      df_f = pd.concat([df_new_row, df_f])

  print(df_f)
  df_f.to_csv(pathCSV +filename+'with_absent.csv', index = False, header=True, sep=';')

mainMenu()
```

**S3 Table** Population abundances (n per 4 m$^2$) and relative genera contributions (%) in our surveys of coral communities in 2021 and 2022.

| 2021 | | | 2022 | | |
|---|---|---|---|---|---|
| **Genus** | **Abundance** | **%** | **Genus** | **Abundance** | **%** |
| *Acropora* | 270 | 30.2 | *Acropora* | 249 | 29.7 |
| *Montipora* | 125 | 14 | *Montipora* | 117 | 14 |
| *Porites* | 108 | 12.1 | *Porites* | 108 | 12.9 |
| *Galaxea* | 78 | 8.7 | *Galaxea* | 71 | 8.5 |
| *Favia* | 50 | 5.6 | *Millepora* | 54 | 6.4 |
| *Millepora* | 47 | 5.3 | *Favia* | 47 | 5.6 |
| *Stylophora* | 41 | 4.6 | *Stylophora* | 31 | 3.7 |
| *Goniastrea* | 26 | 2.9 | *Goniastrea* | 25 | 3 |
| *Pavona* | 23 | 2.6 | *Pavona* | 25 | 3 |
| *Favites* | 21 | 2.3 | *Lobophyllia* | 19 | 2.3 |
| *Lobophyllia* | 20 | 2.2 | *Favites* | 16 | 1.9 |
| *Montastrea* | 15 | 1.7 | *Montastrea* | 15 | 1.8 |
| *Pocillopora* | 14 | 1.6 | *Pocillopora* | 11 | 1.3 |
| *Fungia* | 11 | 1.2 | *Fungia* | 10 | 1.2 |
| *Seriatopora* | 11 | 1.2 | *Seriatopora* | 10 | 1.2 |
| *Leptoseris* | 6 | 0.7 | *Turbinaria* | 5 | 0.6 |
| *Turbinaria* | 5 | 0.6 | *Echinopora* | 4 | 0.5 |
| *Echinopora* | 4 | 0.4 | *Leptoseris* | 4 | 0.5 |
| *Platygyra* | 4 | 0.4 | *Platygyra* | 4 | 0.5 |
| Unidentified | 4 | 0.4 | Unidentified | 4 | 0.5 |
| *Acanthastrea* | 2 | 0.2 | *Acanthastrea* | 2 | 0.2 |
| *Leptoria* | 2 | 0.2 | *Leptoria* | 2 | 0.2 |
| *Pachyseris* | 2 | 0.2 | *Psammocora* | 2 | 0.2 |
| *Psammocora* | 2 | 0.2 | *Astreopora* | 1 | 0.1 |
| *Astreopora* | 1 | 0.1 | *Isopora* | 1 | 0.1 |
| *Cyphastrea* | 1 | 0.1 | *Pachyseris* | 1 | 0.1 |
| *Hydnophora* | 1 | 0.1 | | | |
| **Total** | **894** | **100** | **Total** | **838** | **100** |

**S4 Table** Kolmogorov-Smirnov test results comparing size distributions of the six dominant coral genera in 2021 and 2022. * for *p* <0.05, ** for *p* <0.005, *** for *p* <0.0005, *ns* for non-significant.

| **2021** | | | | | | | |
|---|---|---|---|---|---|---|---|
| | ***Acropora*** | ***Montipora*** | ***Porites*** | ***Galaxea*** | ***Favia*** | **Mean size (cm)** | **Median size (cm)** |
| ***Acropora*** | - | - | - | - | - | 5.7 | 4.5 |
| ***Montipora*** | *ns* | - | - | - | - | 6.3 | 4.9 |
| ***Porites*** | *ns* | *ns* | - | - | - | 6.0 | 4.5 |
| ***Galaxea*** | *** | *** | ** | - | - | 3.8 | 3.6 |
| ***Favia*** | *ns* | * | *ns* | *ns* | - | 5.3 | 3.6 |
| ***Millepora*** | ** | *** | ** | * | *ns* | 4.1 | 2.9 |

| **2022** | | | | | | | |
|---|---|---|---|---|---|---|---|
| | ***Acropora*** | ***Montipora*** | ***Porites*** | ***Galaxea*** | ***Favia*** | **Mean size (cm)** | **Median size (cm)** |
| ***Acropora*** | - | - | - | - | - | 6.4 | 5.5 |
| ***Montipora*** | *ns* | - | - | - | - | 6.1 | 4.7 |
| ***Porites*** | * | *ns* | - | - | - | 5.6 | 4.5 |
| ***Galaxea*** | *** | *** | *** | - | - | 3.4 | 3.2 |
| ***Favia*** | *** | *ns* | *ns* | * | - | 5.1 | 3.6 |
| ***Millepora*** | *** | ** | * | *ns* | *ns* | 4.1 | 3.3 |

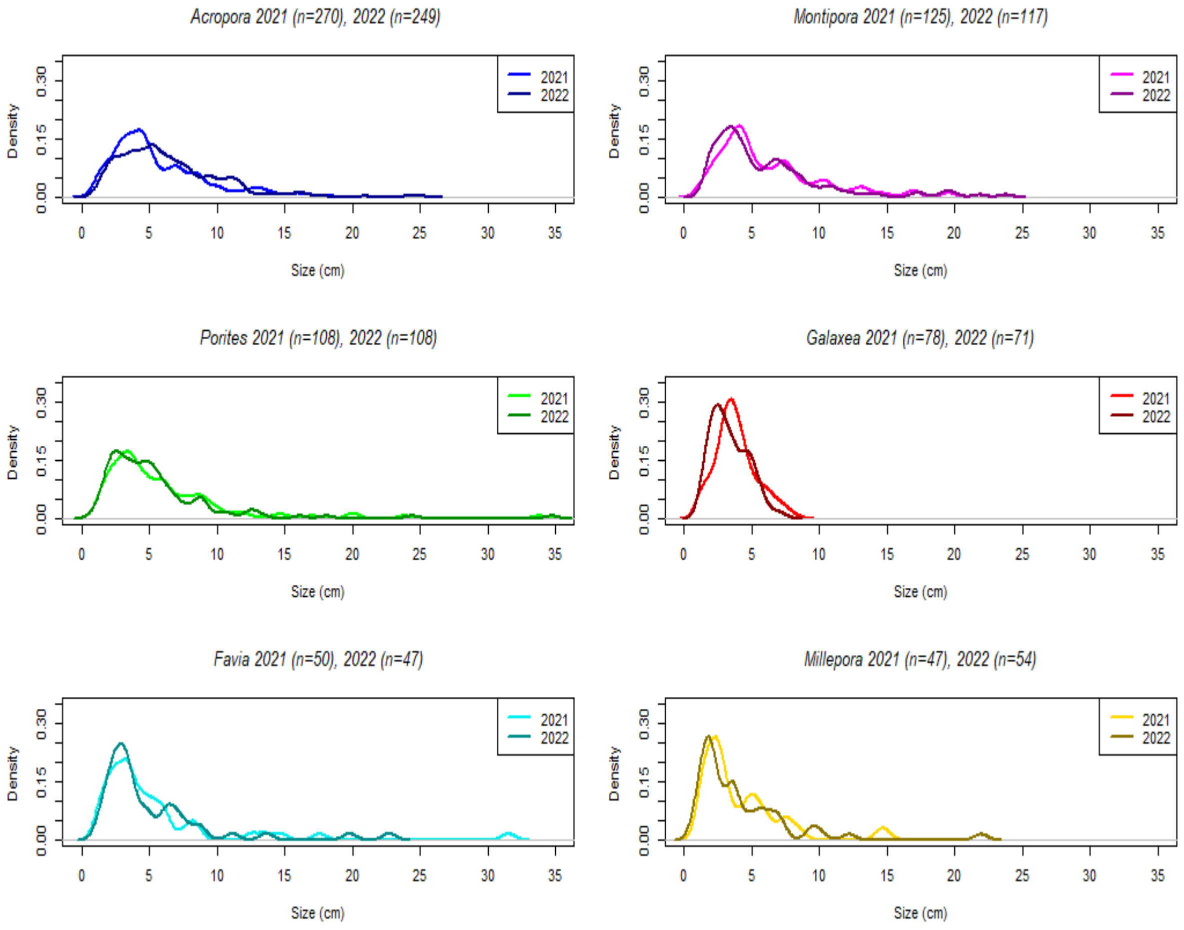


**S5 Figure** Size distributions of populations of the six dominant coral genera in 2021 and 2022.
